## Supplementary Information for "Obligate sexual reproduction of a homothallic fungus closely related to the *Cryptococcus* pathogenic species complex"

### for

#### Supplementary Figures

**A**

| | CBS7841 | CBS7855 |
|---|---|---|
| **Nanopore reads** | | |
| Number of reads | 1,532,981 | 771,560 |
| Total bases | 3,996,171,095 | 2,123,419,399 |
| Mean read length | 2,607 | 2,752 |
| Longest read (bp) | 167,130 | 174,557 |
| Read length N50 | 8,654 | 14,422 |
| **Assembly statistics** | | |
| No. of contigs/chrs. | 8 | 8 |
| Nuclear genome size (bp) | 16,179,376 | 16,328,170 |
| Coverage | 247x | 130x |
| GC% | 44.8 | 44.8 |
| tRNA genes (nuclear) | 98 | 98 |
| Protein-coding genes | 6,342 | 6,329 |
| Telomeric repeats [5'-TAA(C)4-5] | 16 | 16 |

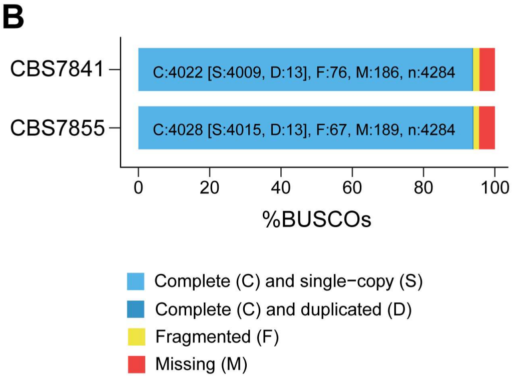


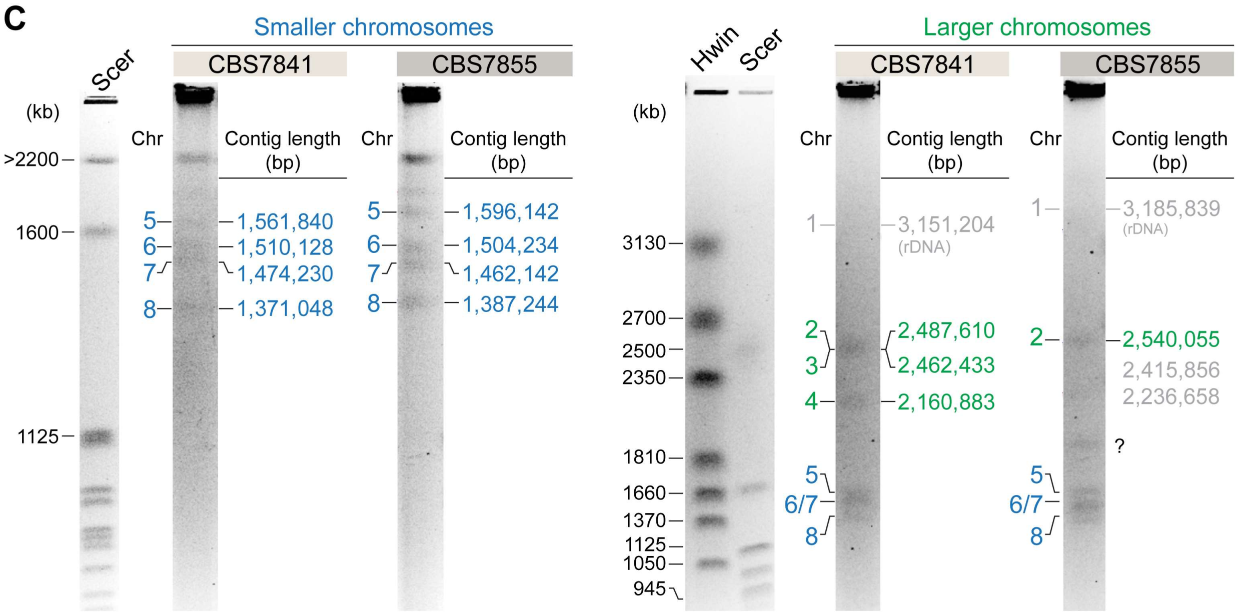


**Figure 1–figure supplement 1**. **(A)** Genome sequencing data generated, final genome assembly statistics, and **(B)** genome completeness assessment upon aligning *C. depauperatus* gene sets to 4284 Benchmarking Universal Single-Copy Orthologs (BUSCOs) included in the tremellomycetes_odb10 dataset. **(C)** Electrophoretic karyotypes (CHEF) of CBS7841 and CBS7855. Chromosomes of *Saccharomyces cerevisiae* (Scer) and *Hansenula wingei* (Hwin) served as size markers. For CBS7841, all contigs (except the largest one that includes the rDNA array, which runs at a higher size range) could be assigned to a band in the gel. For CBS7855, we found two discrepancies: (i) a band (marked with a question mark) that could not be assigned to any of the assembled contigs, and (ii) the fact that contigs 3 and 4 seem to be unaccounted for in the gel. This might suggest inherent chromosome instability in this strain (e.g., a subpopulation of cells undergoing chromosomal rearrangements/breaks/fusions), which will be the subject of future studies.

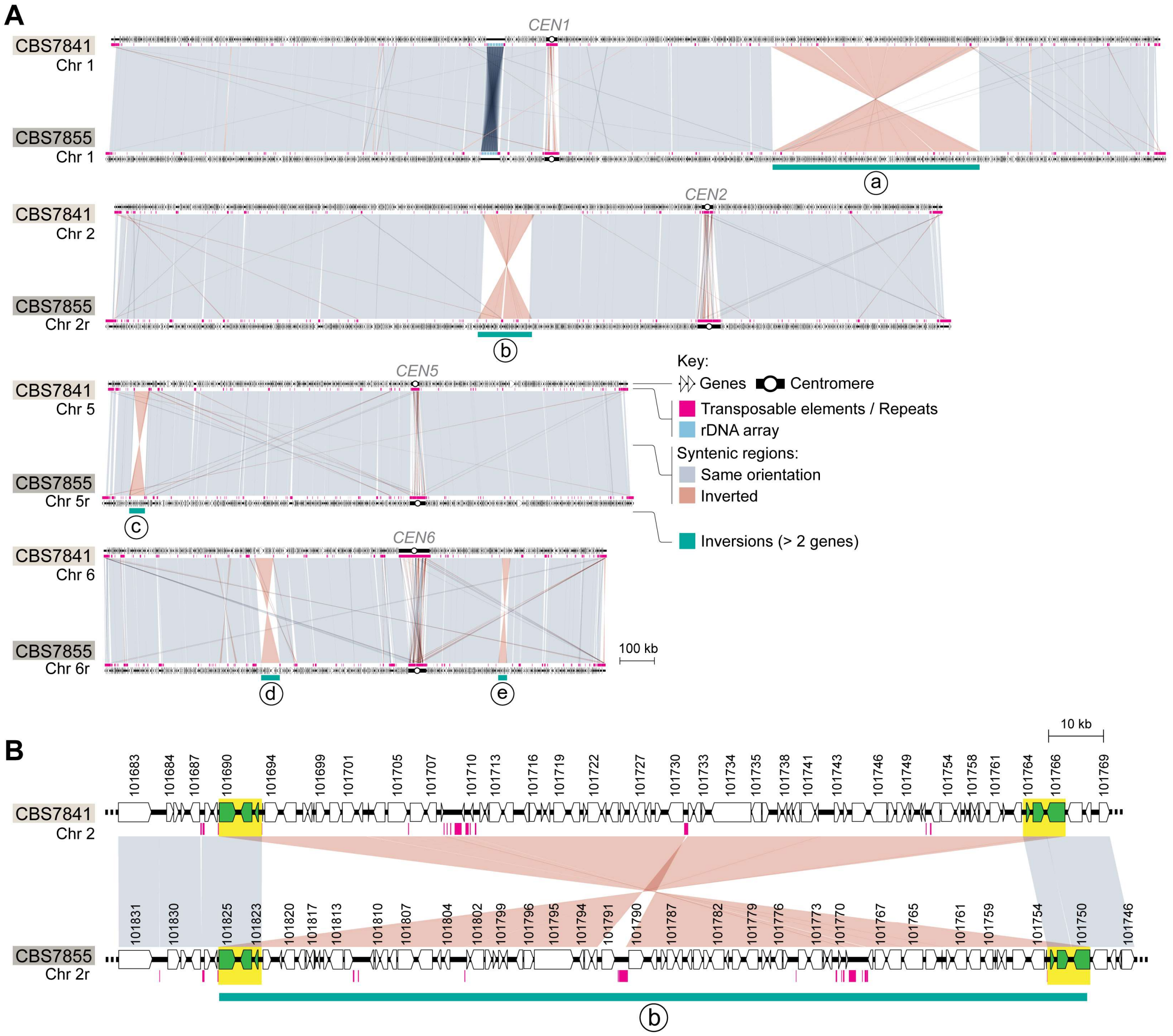


**Figure 1–figure supplement 2. (A)** Synteny maps of *C. depauperatus* CBS7841 and CBS7855 highlighting five large inversions detected between the two strains, with each inversion containing more than two genes (labelled “a” to “e” and denoted by a teal blue bar). **(B)** Except for inversion “a”, all the others (inversion “b” is shown as an example) are associated with duplicated sequences at the borders (highlighted in yellow). Grey or pink bars/lines connect similar regions between the two strains (BLASTN hits > 200 bp) in the same or inverted orientation, respectively. An “r” next to the chromosome number (e.g., Chr 6r in CBS7855) indicates that the orientation is reversed relative to the orientation in the assembly.

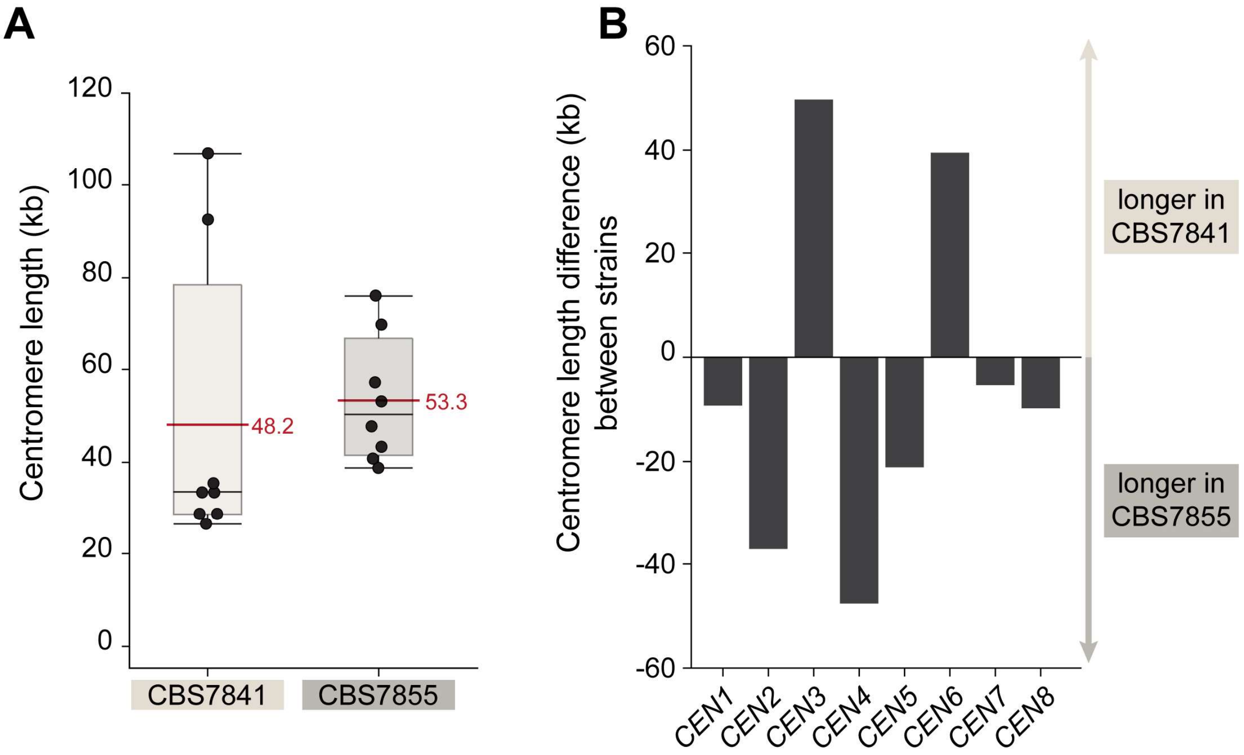


**C**

| | | *CEN* length (bp) | *CEN* coordinates (bp) | | Left *CEN*-flanking gene Gene_ID | coordinates (bp) | | str. | Right *CEN*-flanking gene Gene_ID | coordinates (bp) | | str. |
|---|---|---|---|---|---|---|---|---|---|---|---|---|
| CBS7841 | *CEN1* | 33,986 | 1,306,458 | 1,340,444 | L203_100492 | 1,306,111 | 1,306,457 | + | L203_100493 | 1,340,445 | 1,342,544 | - |
| | *CEN2* | 32,759 | 1,765,455 | 1,798,214 | L203_101986 | 1,763,708 | 1,765,454 | + | L203_101987 | 1,798,215 | 1,799,495 | - |
| | *CEN3* | 106,887 | 366,867 | 473,754 | L203_102433 | 362,798 | 366,866 | + | L203_102434 | 473,755 | 474,974 | - |
| | *CEN4* | 28,508 | 888,831 | 917,339 | L203_103648 | 885,985 | 888,830 | + | L203_103649 | 917,340 | 919,011 | - |
| | *CEN5* | 26,528 | 909,053 | 935,581 | L203_104487 | 905,156 | 909,052 | - | L203_104490 | 935,582 | 937,312 | - |
| | *CEN6* | 92,541 | 885,723 | 978,264 | L203_105096 | 883,838 | 885,722 | + | L203_105097 | 978,265 | 979,451 | - |
| | *CEN7* | 35,365 | 1,078,927 | 1,114,292 | L203_105757 | 1,076,640 | 1,078,926 | + | L203_105758 | 1,114,293 | 1,116,426 | - |
| | *CEN8* | 28,884 | 793,509 | 822,393 | L203_106234 | 792,690 | 793,508 | + | L203_106235 | 822,394 | 823,124 | - |
| CBS7855 | *CEN1* | 43,280 | 1,319,584 | 1,362,864 | L204_100497 | 1,317,052 | 1,319,583 | + | L204_100498 | 1,362,865 | 1,365,213 | - |
| | *CEN2* | 69,763 | 692,075 | 761,838 | L204_101525 | 690,793 | 692,074 | + | L204_101526 | 761,839 | 763,585 | - |
| | *CEN3* | 57,243 | 357,538 | 414,781 | L204_102419 | 353,661 | 357,537 | + | L204_102420 | 414,782 | 416,001 | - |
| | *CEN4* | 76,052 | 1,262,127 | 1,338,179 | L204_103751 | 1,260,458 | 1,262,126 | + | L204_103755 | 1,338,180 | 1,341,026 | - |
| | *CEN5* | 47,715 | 624,967 | 672,682 | L204_104401 | 622,977 | 624,966 | + | L204_104403 | 672,683 | 676,580 | + |
| | *CEN6* | 53,123 | 539,408 | 592,531 | L204_104960 | 538,221 | 539,407 | + | L204_104961 | 592,532 | 594,416 | - |
| | *CEN7* | 40,723 | 1,071,103 | 1,111,826 | L204_105759 | 1,068,052 | 1,071,102 | + | L204_105760 | 1,111,827 | 1,113,960 | - |
| | *CEN8* | 38,698 | 560,745 | 599,443 | L204_106118 | 560,014 | 560,744 | + | L204_106119 | 599,444 | 601,433 | - |

**Figure 1–figure supplement 3.** Centromere length comparison between CBS7841 and CBS7855. **(A)** Box plots depicting the predicted centromere length. Each dot represents one centromere, and the horizontal red lines depict mean centromere lengths in the corresponding strain. Shaded boxes represent the interquartile ranges, and upper or lower whiskers show the largest or smallest observations, respectively. The centromere length differences observed between the two strains are not statistically significant ($P$ = 0.1036; Wilcoxon test). **(B)** Bar graph depicting the length difference between centromeres of homologous chromosomes of the two strains. **(C)** List of ORFs flanking the candidate centromeric regions, genomic coordinates of the predicted centromeres and their exact length. Centromere length was defined as the distance between centromere-flanking ORFs.

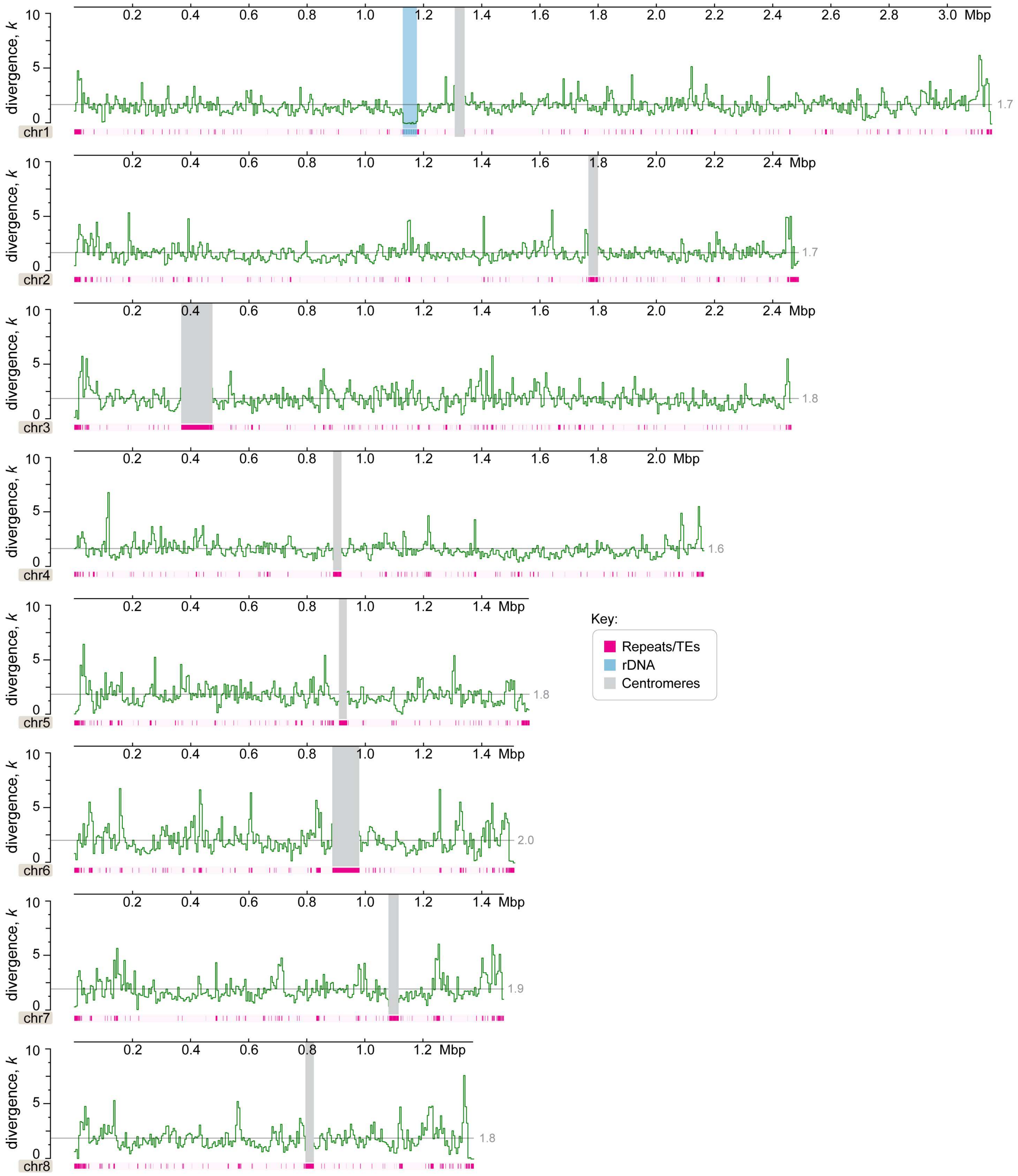


**Figure 1–figure supplement 4.** Genome-wide divergence (*k,* with Jukes-Cantor correction) of *C. depauperatus* CBS7855 relative to CBS7841 genome reference showing no evidence of introgression (which would appear as regions with zero or nearly zero divergence) between the two strains (y-axis values represent percentages; x-axis values represent mega base pairs). Centromeres, rDNA and repeat content are depicted as indicated in the key. Each data point represents the average divergence using a non-overlapping sliding-window of 5,000 sites. A grey line shows the mean divergence for that chromosome.

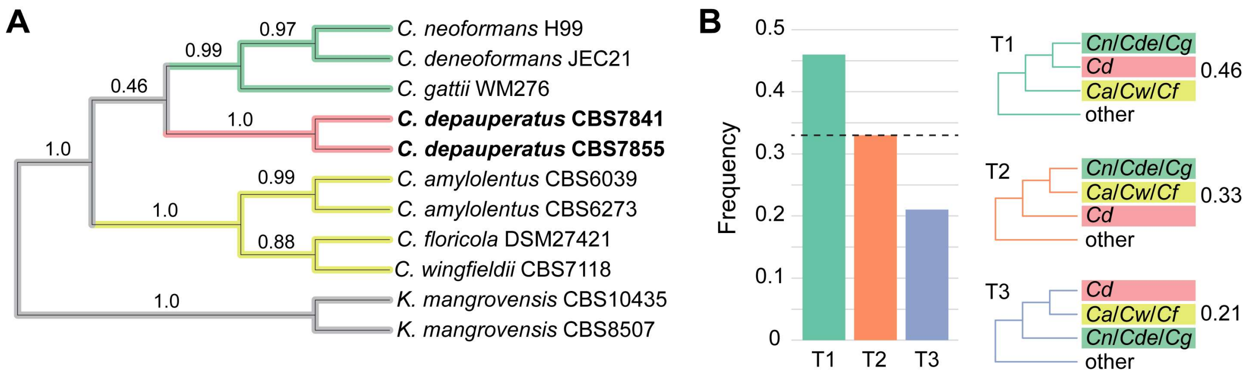


**Figure 1–figure supplement 5. (A)** Tree topology derived from the coalescence approach is fully congruent with the ML tree derived from a concatenation-based approach. Numerical values on the branches represent quartet support for the main topology (q1). All branches received 1.0 local posterior probability (LPP). **(B)** Examination of support among individual gene trees for alternative hypotheses of conflicting branches in the species phylogeny. Green bars and topology (T1) reflect the relationships inferred using a concatenation-based approach on the full data matrix, which also corresponds to the best quartet support, while orange and blue bars and topologies (T2 and T3) correspond to the two alternative hypotheses supported by the two alternative resolutions of each quartet. Dashed horizontal lines mark expectation for a hard polytomy. Abbreviations: *Cn* - *C. neoformans*, *Cde* - *C. deneoformans*, *Cg* - *C. gattii*, *Cd* - *C. depauperatus*, *Ca* - *C. amylolentus*, *Cw* - *C. wingfieldii*, *Cf* - *C. floricola*.

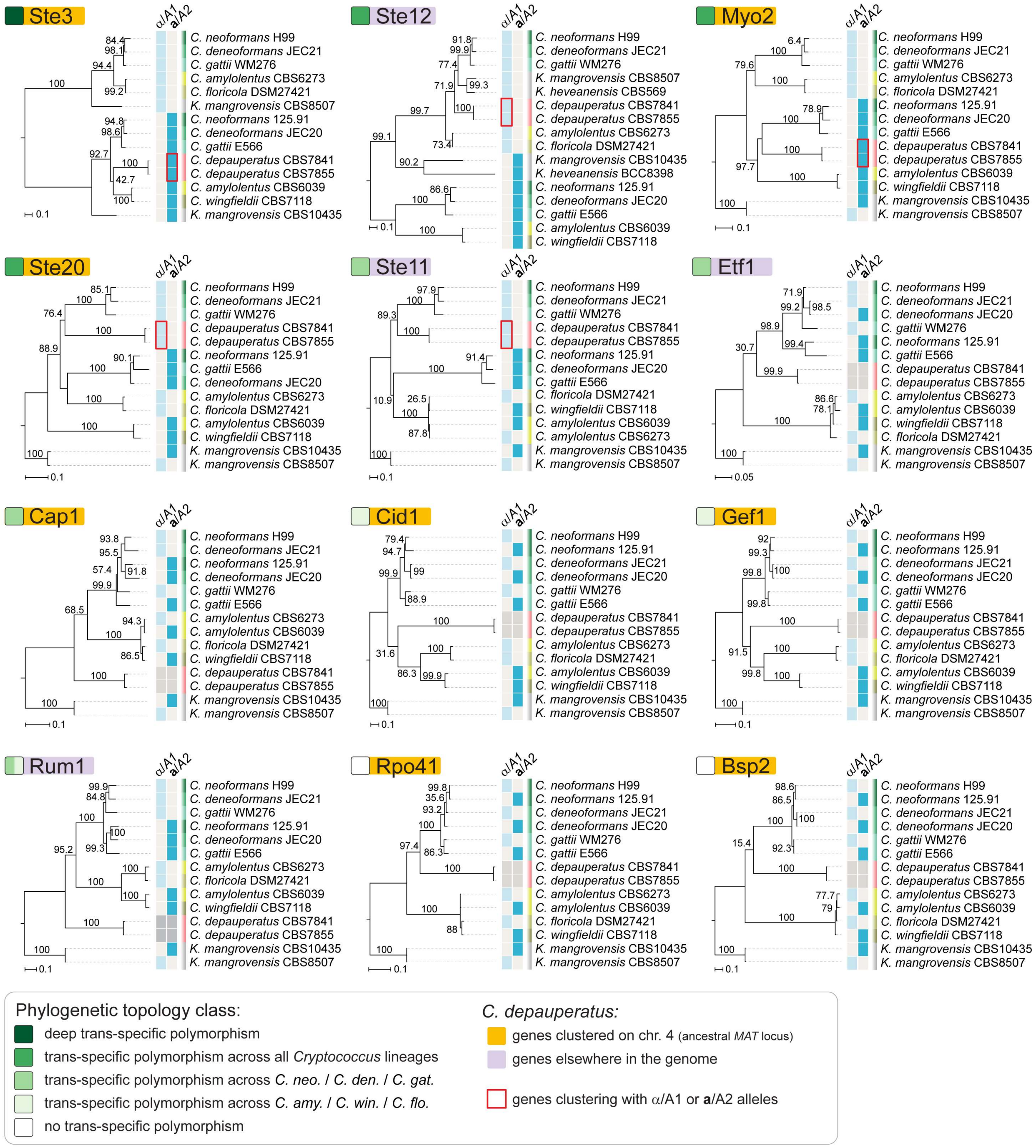


**Figure 2–figure supplement 1.** Individual genealogies of genes within the *MAT* locus of *C. neoformans* exhibiting different level of trans-specific polymorphism**.** Of the analyzed genes, *STE3* and *MYO2* are both found within the predicted ancestral *MAT* region of *C. depauperatus*, and group with strong support with the **a**/A2 alleles from other *Cryptococcus* species. Conversely, *STE12* and *STE11* are found outside the predicted *MAT* region of *C. depauperatus* (each on a different chromosome) and cluster more closely with α/A1 alleles from other *Cryptococcus* species. The *STE20* gene groups more closely with α/A1 alleles despite residing at predicted *MAT* locus in *C. depauperatus*; this may represent the outcome of gene conversion replacing the *STE20***a**/A2 with the α/A1 allele. A deep trans-specific polymorphism is only seen for the *STE3* gene.

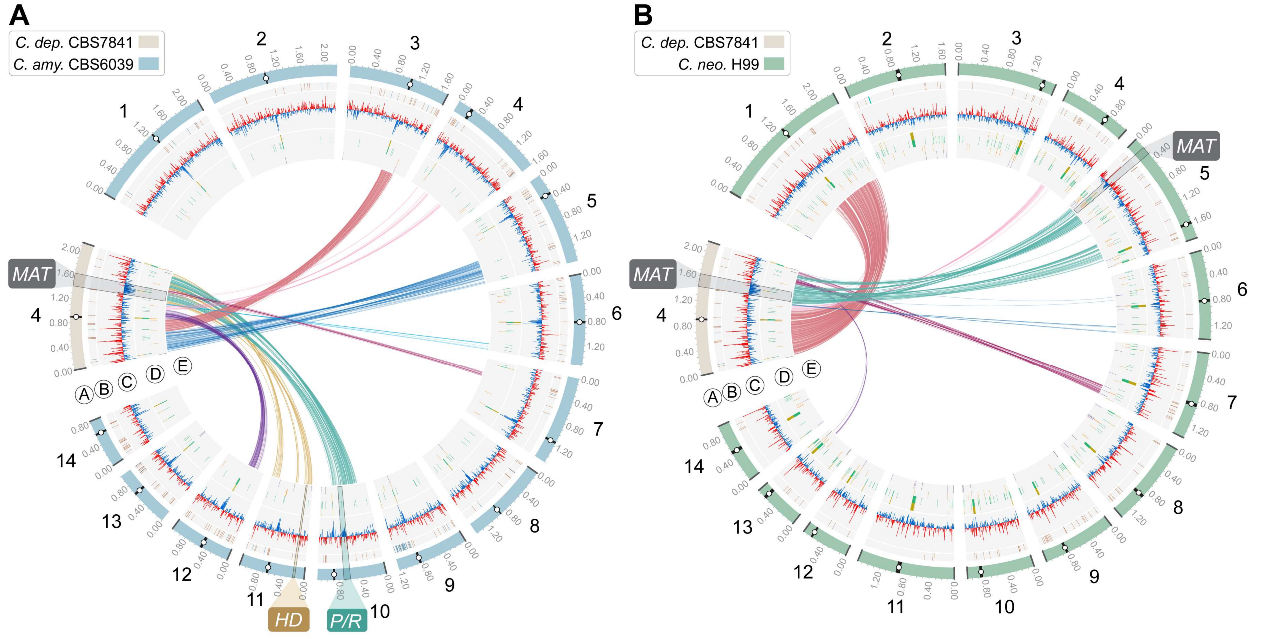


**Figure 2–figure supplement 2.** Genome comparison showing the distribution of BLASTN hits (> 0.5 kb) represented as links between chromosome 4 of *C. depauperatus,* on which the predicted *MAT* locus is located, and all 14 chromosomes of **(A)** *C. amylolentus* CBS6039 and **(B)** *C. neoformans* H99. Tracks "A" to "E" depict genomic features as given in the legend of Figure 1.

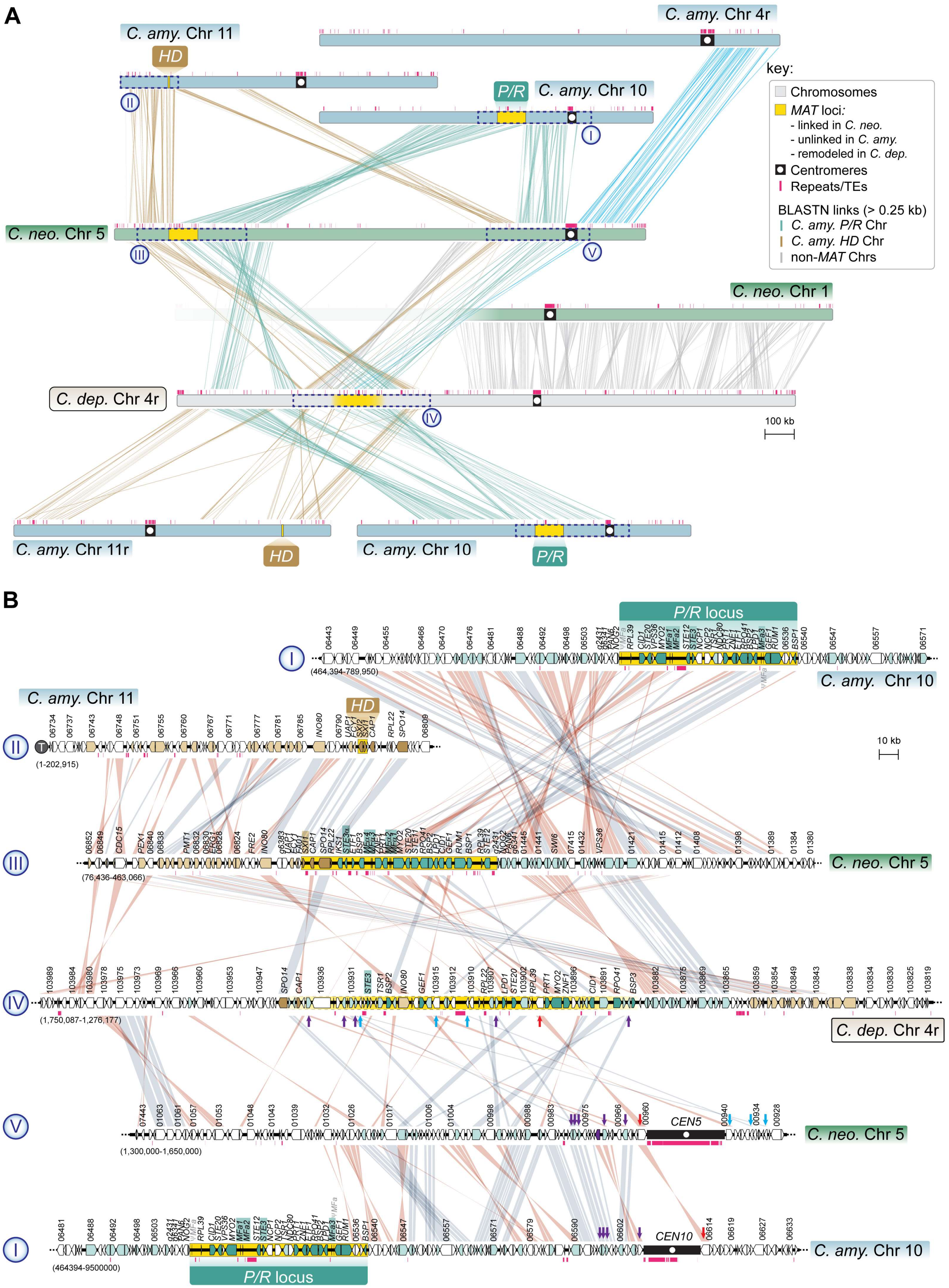
A
C. amy. Chr 4r
C. amy. Chr 11
HD
P/R
C. amy. Chr 10
key:
Chromosomes
MAT loci:
- linked in C. neo.
- unlinked in C. amy.
- remodeled in C. dep.
Centromeres
Repeats/TEs
BLASTN links (> 0.25 kb)
C. amy. P/R Chr
C. amy. HD Chr
non-MAT Chrs
C. neo. Chr 5
C. neo. Chr 1
C. dep. Chr 4r
100 kb
C. amy. Chr 11r
HD
C. amy. Chr 10
P/R
B
P/R locus
C. amy. Chr 10
(464,394-789,950)
C. amy. Chr 11
HD
(1-202,915)
10 kb
C. neo. Chr 5
(76,436-463,066)
C. dep. Chr 4r
(1,750,087-1,276,177)
CEN5
C. neo. Chr 5
(1,300,000-1,650,000)
CEN10
C. amy. Chr 10
(464394-9500000)
P/R locus

**Figure 2–figure supplement 3.** The *MAT* region of *C. depauperatus* contains genes that are flanking the centromeres of *MAT*-containing chromosomes in *C. neoformans* and *C. amylolentus*. **(A)** Linear chromosome plots depicting syntenic regions between the *MAT*-chromosomes of *C. depauperatus* (Chr 4), *C. neoformans* (Chr 5) and *C. amylolentus* (Chrs 10 and 11). In *C. neoformans*, *CEN5* derives from a past event of intercentromeric recombination between *CEN4* and *CEN10* of *C. amylolentus. C. depauperatus CEN4* corresponds to *CEN1* of *C. neoformans*. **(B)** Zoomed-in views of the chromosomal regions enclosed in dashed blue boxes (labelled in roman numerals "I" to "V") showing gene synteny conservation and rearrangements. Note that several genes (denoted by red, purple, and blue arrows) flanking *C. neoformans CEN5* and *C. amylolentus CEN10* are found within the predicted *MAT* region of *C. depauperatus* suggesting this centromere became inactivated following complex chromosomal rearrangements. For simplicity, the locus_tag prefixes (*C. neoformans* H99 - CNAG, *C. amylolentus* CBS6039 - L202, *C. depauperatus* CBS7841 - L203) were omitted from the gene IDs.

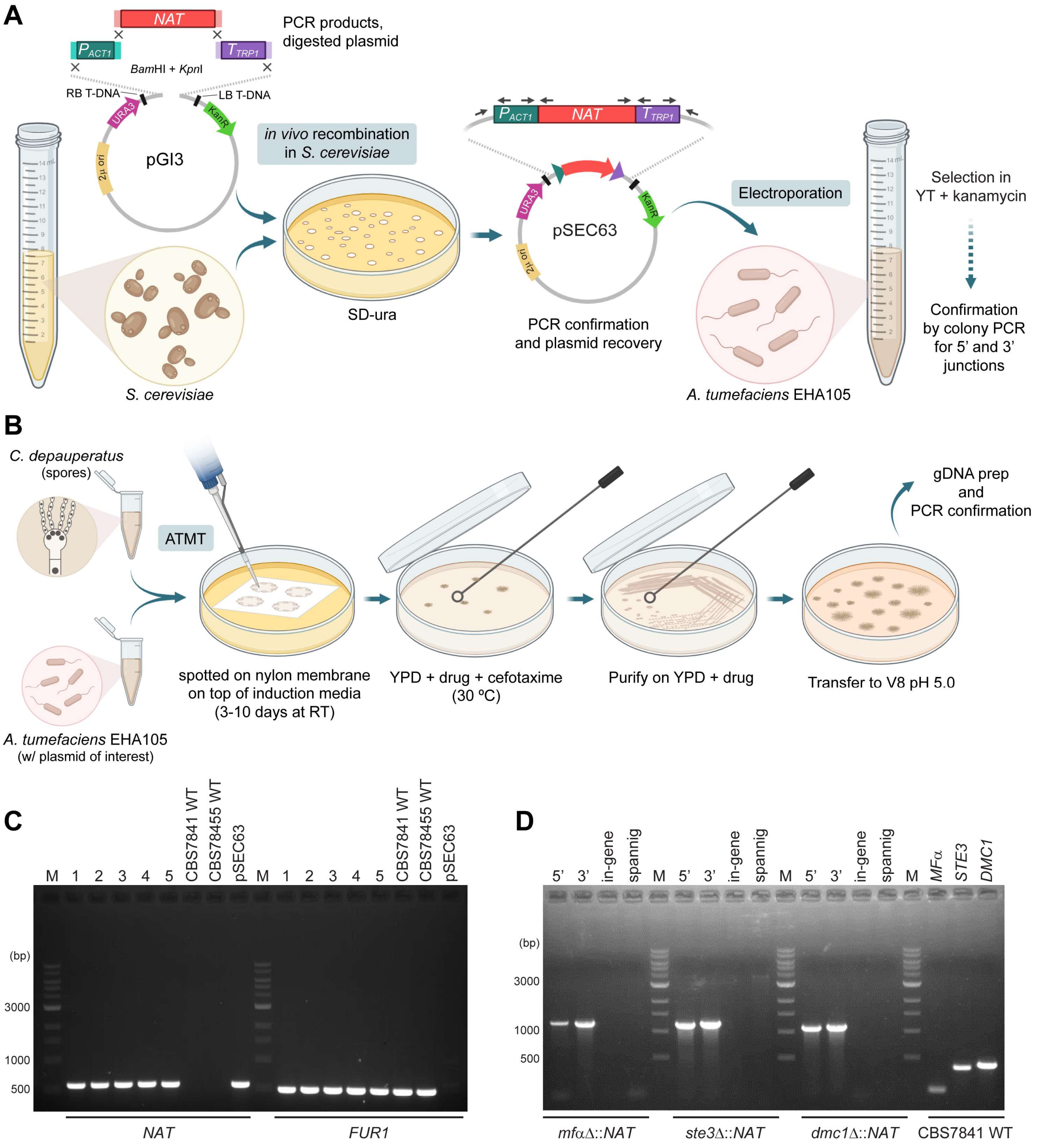


**Figure 5–figure supplement 1. Schematic of the transformation system optimized for *C. depauperatus* and genotypic analysis of *C. depauperatus* transformants**. **(A)** The efficient homologous recombination system of *S. cerevisiae* was employed to construct plasmids for *A. tumefaciens* mediated transformation of *C. depauperatus* as illustrated in panel **(B)**. **(C)** PCR confirmation of ectopic *NAT* transformants. PCR with primers JOHE40162 and JOHE41081 identified the *NAT* gene in CBS7841 (lanes 1-4) and CBS7855 (lane 5) transformants. CBS7841 and CBS7855 wild-type serve as negative controls, and pSEC63 (the plasmid containing the *C. depauperatus* specific *NAT* marker in the binary vector pGI3) serves as a positive control. *FUR1* PCR with primers JOHE45648 and JOHE45649 are positive controls for *C. depauperatus* genomic DNA. **(D)** Junction, in-gene, and spanning PCR confirmation of *mfα*Δ, *ste3*Δ, and *dmc1*Δ mutants. See **Supplementary File 1** for primer sequences. Panels A and B were produced with BioRender.com.

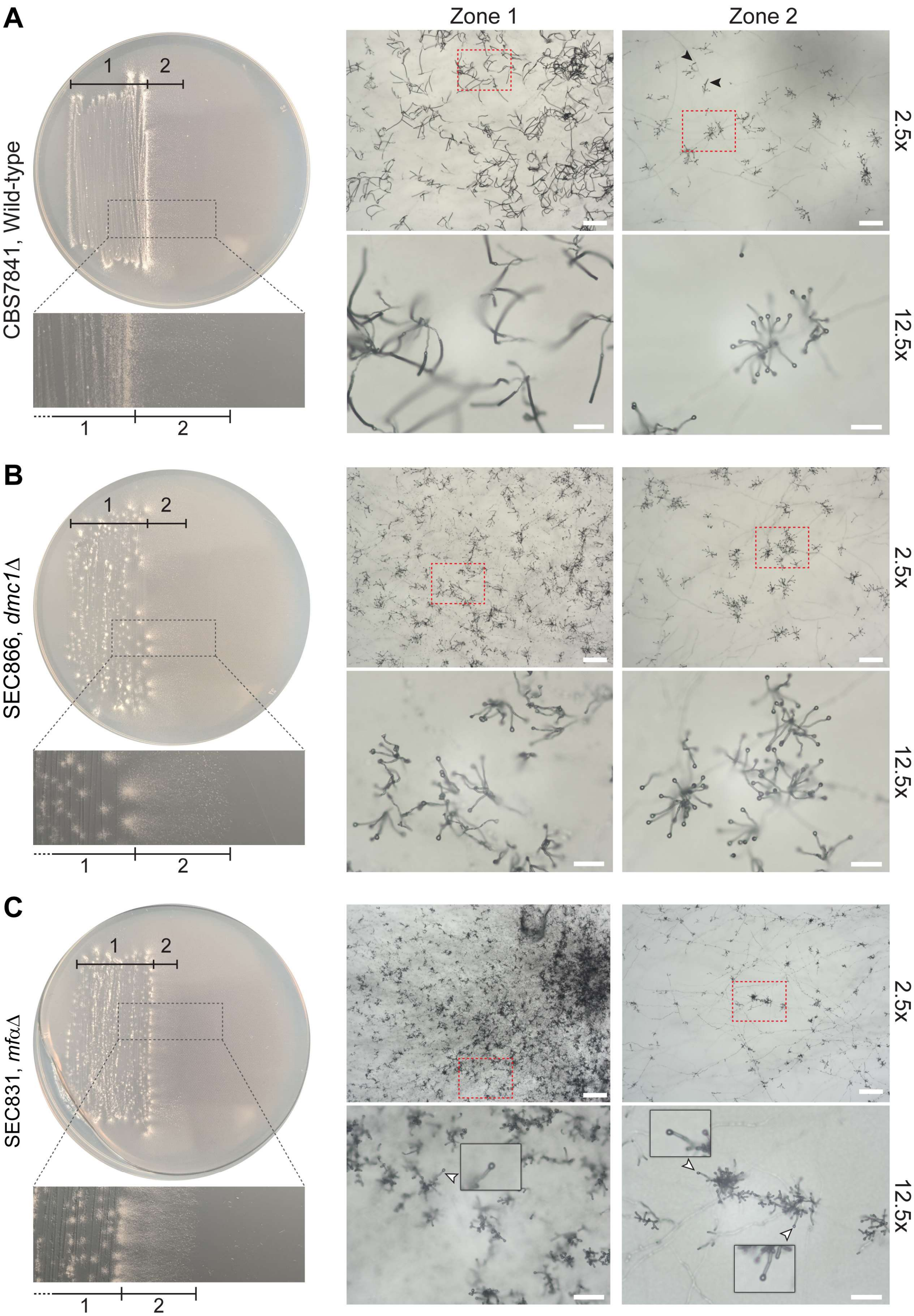
A
B
C
Zone 1
Zone 2
CBS7841, Wild-type
SEC866, dmc1Δ
SEC831, mfαΔ
1
2
2.5x
12.5x

119

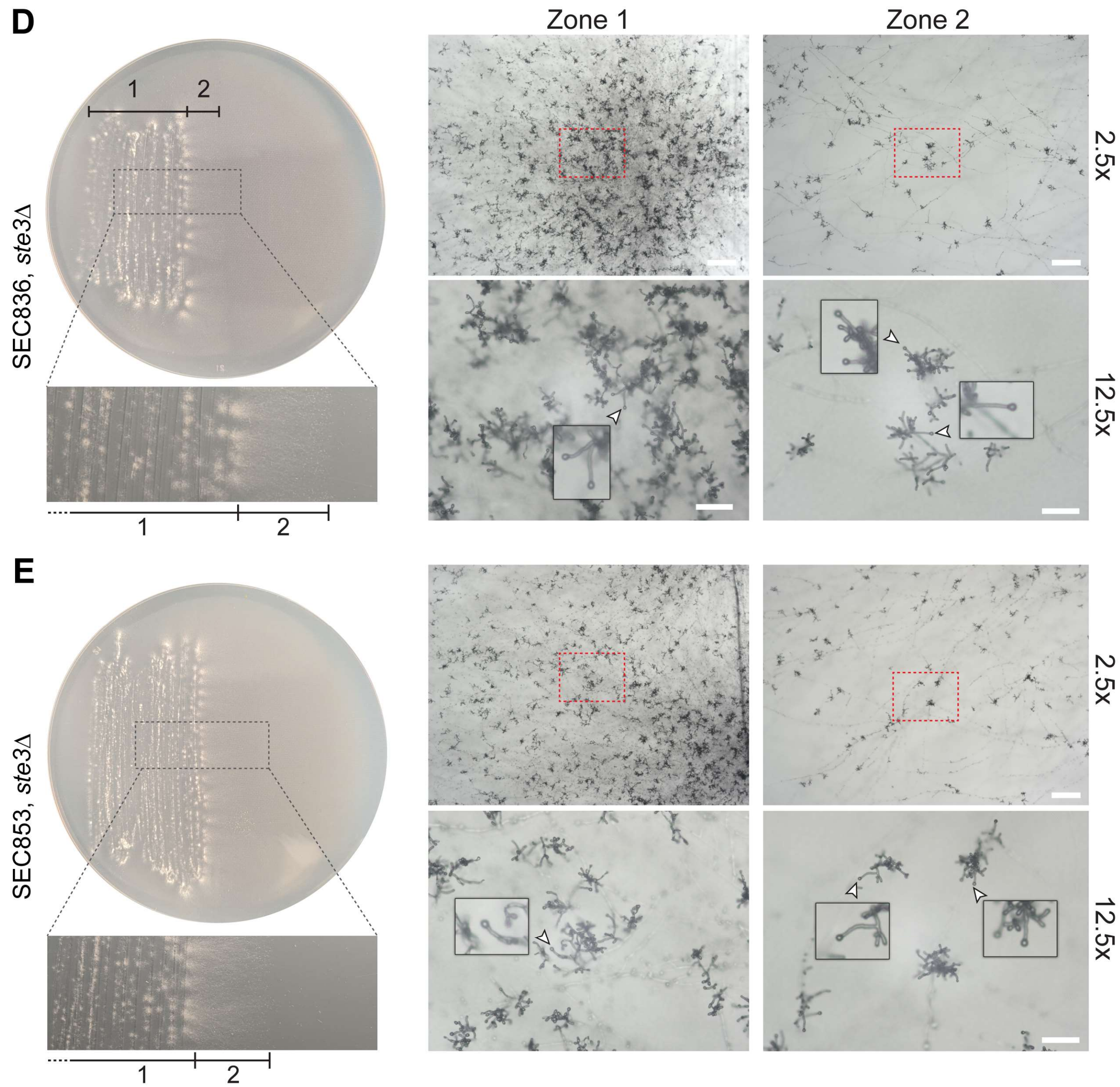


**Figure 5–figure supplement 2**. Representative microscopy images of **(A)** wild-type CBS7841, **(B)** *dmc1*Δ (SEC866), *mf*αΔ (SEC831), **(C)** *ste3*Δ*-1* (SEC836), **(D)**, *ste3*Δ*-2* (SEC853) and **(E)** *dmc1*Δ (SEC866) deletion mutants. Plates were imaged following 25 days of incubation on MS medium at room temperature in the dark. Images were taken at 2.5x (bars = 200 μm) and 12.5x magnification (bars = 50 μm) from two different zones of the culture: zone 1 corresponding to older hyphae (in the wild-type, this zone predominantly contains sporulated basidia) and zone 2 corresponding to younger, actively growing hyphae (in the wild type, this zone predominantly contains unsporulated basidia; a few sporulating basidia are indicated by black arrowheads). Basidia diameter was measured with ImageJ from images taken at 12.5x magnification (see **Figure 5–source data 2**). For each representative image taken at 2.5x magnification, a zoomed-in view (12.5x) corresponding to the region indicated by dashed red squares is shown below. For the *mf*αΔ (SEC831), *ste3*Δ*-1* (SEC836) and *ste3*Δ*-2* (SEC853) mutants, examples of immature basidia are indicated by white arrowheads and shown as zoomed-in insets.

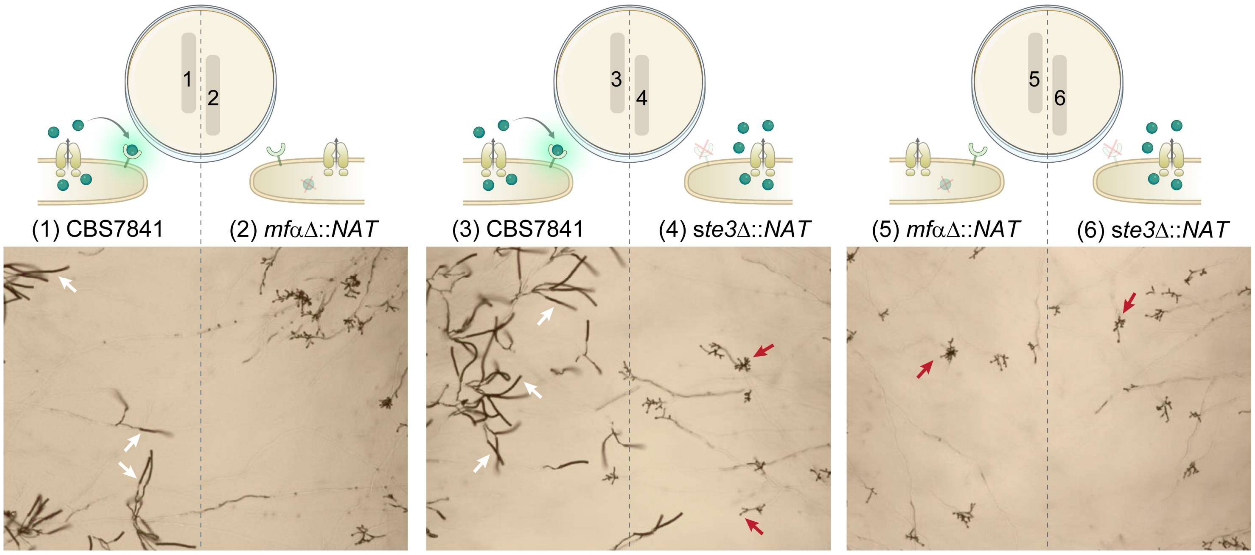


**Figure 6–figure supplement 1. *C. depauperatus* mutant strains do not undergo sporulation in confrontation assays.** Spores and/or hyphae from wild-type and mutant strains were struck ~2 mm apart on MS medium and incubated at room temperature for 2 weeks. Sporulating basidia (a few examples emphasized by white arrows) are observed on the left side of the plate (1 and 3) where the WT strain was inoculated, but not on the right side of the plate corresponding to the mutant strains (red arrows indicate a few examples of bald basidia). The lack of complementation of the *mfαΔ* mutation (expected to occur in 2 and 5) might be due to low diffusion and/or concentration of the mating pheromone, preventing long distance interactions.

**Figure 7–figure supplement 1.** Sanger-sequencing of *CAN1* and *FUR1* loci from double-drug resistant isolates recovered from *C. depauperatus* intra- and inter-strain crosses. Isolates marked in green did not inherit both parental mutant alleles; rather, they gained de novo spontaneous mutations in *CAN1* or *FUR1* loci, or in other genes.

| | Parent 1 | Parent 2 | Double-drug resistant isolates | | |
|---|---|---|---|---|---|
| | | | **#** | ***CAN1* mutation** | ***FUR1* mutation** |
| Intra-strain mating | CBS7841 *can1-2* del1868 | CBS7841 *fur1-1* A774G | 1 | *can1-2* del1868 | *fur1-1* A774G |
| | | | 2 | *can1-2* del1868 | *fur1-1* A774G |
| | | | 3 | *can1-2* del1868 | *fur1-1* A774G |
| | | | 4 | *can1-2* del1868 | *fur1-1* A774G |
| | | | 5 | *can1-2* del1868 | no *FUR1* mutation identified* |
| | | | 6 | *can1-2* del1868 | *fur1-1* A774G |
| | CBS7855 *can1-3* G456A | CBS7855 *fur1-2* G1383T | 1 | *can1-3* G456A | *fur1-2* G1383T |
| | | | 2 | *can1-3* G456A | *fur1 G727A* |
| | | | 3 | *can1-3* G456A | *fur1-2* G1383T |
| | | | 4 | *can1* T611C | *fur1-2* G1383T |
| | | | 5 | *can1* del1488TCT | *fur1-2* G1383T |
| | | | 6 | *can1* del1488TCT | *fur1-2* G1383T |
| | | | 7 | *can1* T611C | *fur1-2* G1383T |
| | | | 8 | *can1-3* G456A | no *FUR1* mutation identified* |
| Inter-strain mating | CBS7855 *can1-3* G456A | CBS7841 *fur1-1* A774G | 1 | *can1* G1899T | *fur1-1* A774G |
| | | | 2 | *can1* A39G | *fur1-1* A774G |
| | | | 3 | *can1* C367T | *fur1-1* A774G |
| | | | 4 | *can1* G1423A | *fur1-1* A774G |
| | | | 5 | *can1* G1423A | *fur1-1* A774G |
| | | | 6 | *can1-3* G456A | *fur1* G727A |
| | | | 7 | *can1-3* G456A | no *FUR1* mutation identified* |
| | | | 8 | *can1-3* G456A | no *FUR1* mutation identified* |
| | CBS7841 *can1-2* del1868 | CBS7855 *fur1-2* G1383T | 1 | *can1-2* del1868 | no *FUR1* mutation identified* |
| | | | 2 | *can1* T611C | *fur1-2* G1383T |
| | | | 3 | *can1-2* del1868 | no *FUR1* mutation identified* |
| | | | 4 | *can1-2* del1868 | *fur1* A774G |

*In these cases, resistance to 5-FC is likely conferred by mutations in other genes, including *FCY1*, *FCY2*, *UXS1*, or *TCO2*.

#### Supplementary Files

**Supplementary File 1.** Strains used in this study.

**Supplementary File 2.** Primers used in this study.
