## Supplementary material for "Obligate sexual reproduction of a homothallic fungus closely related to the *Cryptococcus* pathogenic species complex": Figure 5-source data 1

**Figure 5–source data 1.** Frequency of basidia defective in sporulation (bald basidia) and basidia with spores in *C. depauperatus* wild-type (CBS7841) and mutant strains, and one-way ANOVA and Tukey's HSD post hoc statistical tests for frequencies of bald basidia.

| **Strain** | **Genotype** | **Areas surveyed** | **Bald basidia** | **Basidia w/ spores** | **Total basidia** | **Frequency of bald basidia (%)** | **Frequency of basidia with spores (%)** |
|---|---|---|---|---|---|---|---|
| CBS7841 | WT | 1 | 0 | 25 | 25 | 0 | 100 |
| | | 2 | 1 | 35 | 36 | 2.78 | 97.22 |
| | | 3 | 1 | 38 | 39 | 2.56 | 97.44 |
| SEC831 | *mfα∆* | 1 | 16 | 3 | 19 | 84.21 | 15.79 |
| | | 2 | 15 | 3 | 18 | 83.33 | 16.67 |
| | | 3 | 11 | 0 | 11 | 100 | 0 |
| | | 4 | 10 | 0 | 10 | 100 | 0 |
| | | 5 | 15 | 0 | 15 | 100 | 0 |
| | | 6 | 18 | 0 | 18 | 100 | 0 |
| SEC836 | *ste3∆* | 1 | 18 | 0 | 18 | 100 | 0 |
| | | 2 | 5 | 0 | 5 | 100 | 0 |
| | | 3 | 17 | 0 | 17 | 100 | 0 |
| | | 4 | 16 | 0 | 16 | 100 | 0 |
| | | 5 | 17 | 5 | 22 | 77.27 | 22.73 |
| SEC866 | *dmc1∆* | 1 | 39 | 0 | 39 | 100 | 0 |
| | | 2 | 42 | 0 | 42 | 100 | 0 |
| | | 3 | 19 | 0 | 19 | 100 | 0 |
| #B6.1 | *mfa∆ ste3∆* | 1 | 11 | 0 | 11 | 100 | 0 |
| | | 2 | 23 | 0 | 23 | 100 | 0 |
| | | 3 | 29 | 0 | 29 | 100 | 0 |
| | | 4 | 5 | 0 | 5 | 100 | 0 |
| | | 5 | 32 | 0 | 32 | 100 | 0 |

### One-way ANOVA

| Source | Degrees of freedom | Sum of Squares | Mean Square | F Ratio | p-value |
|---|---|---|---|---|---|
| Cross type | 4 | 23656.868 | 5914.22 | 130.6550 | **<.0001*** ** |
| Residuals | 17 | 769.520 | 45.27 | | |

### Tukey's HSD

| | | | 95% Confidence interval | | |
|---|---|---|---|---|---|
| Level | - Level | Difference | Lower | Upper | p-Value |
| CBS7841 | *dmc1∆* | 98.21937 | 81.5059 | 114.9329 | **<.0001*** |
| CBS7841 | *mfα∆ ste3∆* #B6.1 | 98.21937 | 83.2704 | 113.1684 | **<.0001*** |
| CBS7841 | *ste3∆* | 93.67392 | 78.7249 | 108.6229 | **<.0001*** |
| CBS7841 | *mfα∆* | 92.81002 | 78.3357 | 107.2843 | **<.0001*** |
| *mfα∆* | *dmc1∆* | 5.40936 | -9.0650 | 19.8837 | 0.7852 |
| *mfα∆* | *mfα∆ ste3∆* #B6.1 | 5.40936 | -6.9857 | 17.8044 | 0.6786 |
| *ste3∆* | *dmc1∆* | 4.54545 | -10.4036 | 19.4945 | 0.8833 |
| *ste3∆* | *mfα∆ ste3∆* #B6.1 | 4.54545 | -8.4008 | 17.4917 | 0.8199 |
| *mfα∆* | *ste3∆* | 0.86390 | -11.5312 | 13.2590 | 0.9995 |
| *mfα∆ ste3∆* #B6.1 | *dmc1∆* | 0.00000 | -14.9490 | 14.9490 | 1.0000 |
