## Supplementary material for "Obligate sexual reproduction of a homothallic fungus closely related to the *Cryptococcus* pathogenic species complex": Figure 6-source data 1

**Figure 6–source data 1.** Frequency of basidia defective in sporulation (bald basidia) and basidia with spores in *C. depauperatus* wild-type (CBS7841), *mf*α∆ and *ste3*∆ single mutants, and the *mf*α∆ x *ste3*α∆ co-cultures mutant strains and one-way ANOVA and Tukey's HSD post hoc statistical tests for frequencies of sporulating basidia. Related to Figure 6B.

| **Strain** | **Genotype** | **Areas surveyed** | **Bald basidia** | **Basidia w/ spores** | **Total basidia** | **Frequency of bald basidia (%)** | **Frequency of basidia with spores (%)** |
|---|---|---|---|---|---|---|---|
| CBS7841 | WT | 1 | 2 | 23 | 25 | 8 | 92 |
| | | 2 | 0 | 26 | 26 | 0 | 100 |
| | | 3 | 4 | 26 | 30 | 13.33 | 86.67 |
| | | 4 | 0 | 15 | 15 | 0 | 100 |
| SEC831 | *mf*α∆ | 1 | 37 | 0 | 37 | 100 | 0 |
| | | 2 | 33 | 1 | 34 | 97.06 | 2.94 |
| | | 3 | 29 | 0 | 29 | 100 | 0 |
| SEC836 | *ste3*∆ | 1 | 18 | 0 | 18 | 100 | 0 |
| | | 2 | 18 | 0 | 18 | 100 | 0 |
| | | 3 | 22 | 0 | 22 | 100 | 0 |
| | | 4 | 39 | 0 | 39 | 100 | 0 |
| SEC831 x SEC836 | *mfa*∆ x *ste3*∆ | 1 | 15 | 1 | 16 | 93.75 | 6.25 |
| | | 2 | 19 | 1 | 20 | 95 | 5 |
| | | 3 | 20 | 4 | 24 | 83.33 | 16.67 |
| | | 4 | 25 | 3 | 28 | 89.29 | 10.71 |
| | | 5 | 8 | 4 | 12 | 66.67 | 33.33 |

### One-way ANOVA

| Source | Degrees of freedom | Sum of Squares | Mean Square | F Ratio | p-value |
|---|---|---|---|---|---|
| Cross type | 3 | 24028.413 | 8009.47 | 144.4004 | **<.0001*** ** |
| Residuals | 12 | 665.605 | 55.47 | | |

### Tukey's HSD

| | | | 95% Confidence interval | | |
|---|---|---|---|---|---|
| Level | - Level | Difference | Lower | Upper | p-Value |
| CBS7841 | SEC836 | 94.66750 | 83.1933 | 106.1417 | **<.0001*** ** |
| CBS7841 | SEC831 | 93.68750 | 81.2939 | 106.0811 | **<.0001*** ** |
| CBS7841 | SEC831 x SEC836 | 80.27550 | 69.3901 | 91.1609 | **<.0001*** ** |
| SEC831 x SEC836 | SEC836 | 14.39200 | 3.5066 | 25.2774 | **0.0138*** |
| SEC831 x SEC836 | SEC831 | 13.41200 | 1.5615 | 25.2625 | **0.0297*** |
| SEC831 | SEC836 | 0.98000 | -11.4136 | 13.3736 | 0.8661 |
